# Molecular basis for a novel systemic form of human hereditary apoA-I amyloidosis with vision loss

**DOI:** 10.1101/354001

**Authors:** Isabel Morgado, Pierre-Raphael Rothschild, Afra Panahi, Jean-Claude Aldigier, Andrew G. Burwash, Madhurima Das, Magali Colombat, Thierry Frouget, Jean Philippe Rerolle, François Paraf, Nathalie Rioux-Leclercq, Jean-Michel Goujon, Caroline Beugnet, Antoine Durrbach, Didier Samuel, Antoine Brézin, John E. Straub, Olga Gursky, Sophie Valleix

**Author notes:** I. Morgado, P.-R. Rothschild and A. Panahi contributed equally to this work. J.-C. Aldigier, A. G. Burwash and M. Das contributed equally to this work. O. Gursky, and S. Valleix contributed equally to this work. **Subject categories:** Human Biology and Medicine; Genetics; Biophysics and Structural Biology.

## Abstract

Hereditary apolipoprotein A-I (apoA-I) amyloidosis (AApoAI) is a life-threatening incurable genetic disorder whose molecular underpinnings and the full spectrum of afflicted organs are unclear. We report a new form of AApoAI with amyloid deposition in multiple organs, including an unprecedented retinal amyloidosis. Genetic and proteomic analyses identified Glu34Lys apoA-I as the fibrillar protein causing the clinical manifestations. A life-saving combined hepatorenal transplantation was performed for one Glu34Lys carrier. To elucidate structural underpinnings for amyloidogenic properties of Glu34Lys, we generated its recombinant globular domain and compared the conformation and dynamics of its lipid-free form with those of two other naturally occurring apoA-I variants, Phe71Tyr (amyloidogenic) and Leu159Arg (non-amyloidogenic). All variants showed reduced stability and altered aromatic residue packing. Molecular dynamics simulations revealed local helical unfolding and suggested that transient opening of Trp72 induced mutation-dependent structural perturbations in a sensitive region, including the major amyloid hotspot residues 14-22. We posit that a shift from the “closed” to an “open” orientation of Trp72 modulates structural protection of amyloid hotspots, suggesting a previously unknown early step in protein misfolding.

## Introduction

Amyloid diseases, including neurodegenerative and systemic amyloidoses, affect millions of patients worldwide and are a major public health challenge. The hallmark of these diverse diseases is misfolding of a protein, from its native structure into an abnormal intermolecular cross-β-sheet, leading to deposition of insoluble amyloid fibrils in organs and tissues.

Apolipoproteins (apos) are a family of lipid surface-binding proteins overrepresented in acquired and hereditary amyloidosis (Nichols *et al*, 1988; Teoh *et al*, 2011; Das & Gursky, 2015). ApoA-I (243 amino acids) is the major structural and functional constituent of plasma HDL that transports cholesterol from peripheral cells and protects against cardiovascular disease and inflammation (Rosenson *et al*, 2012). Numerous naturally occurring *APOA1* gene mutations lead to reduced plasma levels of HDL and apoA-I; some mutations, such as Leu159Arg, are associated with abnormal lipid metabolism and increased risk of atherosclerosis, while others cause amyloidosis *in vivo* (Sorci-Thomas *et al*, 2012). Most circulating apoA-I is stabilized by binding to HDL, yet a fraction of apoA-I is released from HDL in a labile free state that forms the protein precursor of amyloid (Teoh *et al*, 2011; Das & Gursky, 2015; Jayaraman *et al*, 2017). Human apoA-I contains two structural domains, a globular N-terminal domain (residues 1-184) and a flexible hydrophobic C-terminal tail (residues 185-243). The x-ray crystal structure of the globular domain, which forms a four-helix bundle in solution, is the only currently available high-resolution structure of lipid-free apoA-I (Mei *et al*, 2016).

Hereditary apoA-I amyloidosis (AApoAI) is a rare monogenic disease wherein specific mutations in the *APOA1* gene cause abnormal amyloid deposition of predominantly N-terminal fragments of the variant protein in various organs, leading to organ damage (Nichols *et al*, 1988; Obici *et al*, 1999). The available treatments for this debilitating life-threatening disease are limited to haemodialysis and organ transplant (Gillmore *et al*, 2006). Approximately 20 AApoAI variants have been reported worldwide (http://www.amyloidosismutations.com), with mutation sites located in two regions of the globular domain. “Inside” mutations cluster in residues 26-107 from N-terminal 9-11 kDa segments that form fibrillar deposits in AApoAI, while “outside” mutations cluster in residues 154-178 (Obici *et al*, 2006; Gursky *et al*, 2012). No amyloidogenic mutations have been identified in the C-terminal tail, suggesting that it does not directly initiate the misfolding. Structural studies have postulated that AApoAI mutations perturb the native packing in the globular domain, thereby increasing solvent exposure of the major amyloid “hotspot” residues 14-22; increased exposure of this adhesive segment was proposed to trigger amyloid formation ((Das *et al*, 2014; Das *et al*, 2016) and references therein). The exact molecular events whereby amyloidogenic mutations trigger apoA-I misfolding remain unknown. Identifying such early events in the misfolding pathway may help develop much-needed therapeutic targets for this hitherto incurable disease.

AApoAI is an autosomal dominant disorder, yet sporadic cases with no family history have also been reported, complicating the diagnosis (Rowczenio *et al*, 2011). Whether sporadic cases reflect variable penetrance of the disease or result from *de novo APOA1* gene mutations is unknown. Also intriguing is the diverse clinical presentation of the disease. AApoAI can have an early onset with an extensive systemic amyloid deposition in multiple organs (mainly the kidney, liver and heart) leading to a fatal outcome in the fourth to fifth decade of life; alternatively, AApoAI may have a mild phenotype restricted to amyloid in only one organ, usually the oropharyngeal tractus or skin. The molecular basis for this diverse clinical presentation is unclear and probably stems, in part, from distinct properties of apoA-I variants.

Here, we show that Glu34Lys apoA-I variant is the fibril-forming protein that caused a new form of a severe systemic AApoAI, including unprecedented massive amyloid deposits in the retina. To elucidate structural underpinnings for the amyloidogenic properties of Glu34Lys apoA-I, we generated the globular domain of this human variant and explored its structure, stability and dynamics using circular dichroism (CD) spectroscopy and molecular dynamics (MD) simulations. Our results provide new insights into the early steps of the protein misfolding pathway.

## Results

### Clinical studies of two unrelated kindreds show amyloid deposits in multiple organs

Family A (Fig 1A): A 23 year-old man (proband III.3) presented with an eight-year history of hypertension, proteinuria (4.5 g / 24 h), and renal dysfunction (serum creatinine at 2.1 mg/dl) as starting symptoms, followed by infertility and transient episodes of bilateral visual loss and metamorphopsia. Ultrasound of the abdomen revealed hepatomegaly and splenomegaly; electrocardiogram and echocardiography did not reveal any evidence of cardiomyopathy. There was no history of paraesthesia or autonomic symptoms, and no evidence of a plasma cell dyscrasia. Baseline tests showed abnormal renal and liver functions (alkaline phosphatases: 571 U/l, aspartate aminotransferase: 137 U/l, alanine aminotransferase: 88 U/l, gamma-glutamyltransferase: 994 U/l). Plasma triglycerides, total cholesterol, HDL-cholesterol, LDL-cholesterol, and the plasma levels of apoA-I and apoB were in the normal range. Endocrine testing demonstrated hypergonadotrophic hypogonadism with a dramatic drop in plasma concentrations of 17β-estradiol, testosterone, and dihydrotestosterone, and increased plasma concentrations of follicle-stimulating hormone and luteinizing hormone. A kidney biopsy revealed large amounts of eosinophilic Congo-Red positive amyloid deposits in the glomerular compartment (Fig 1C). Amyloid deposits were also found in the liver (Fig 1D). Testicular histology showed massive amyloid deposits in the interstitial tissue associated with a severely reduced number of seminiferous tubules without spermatogenesis (Fig 1F and G).

**Figure 1.**
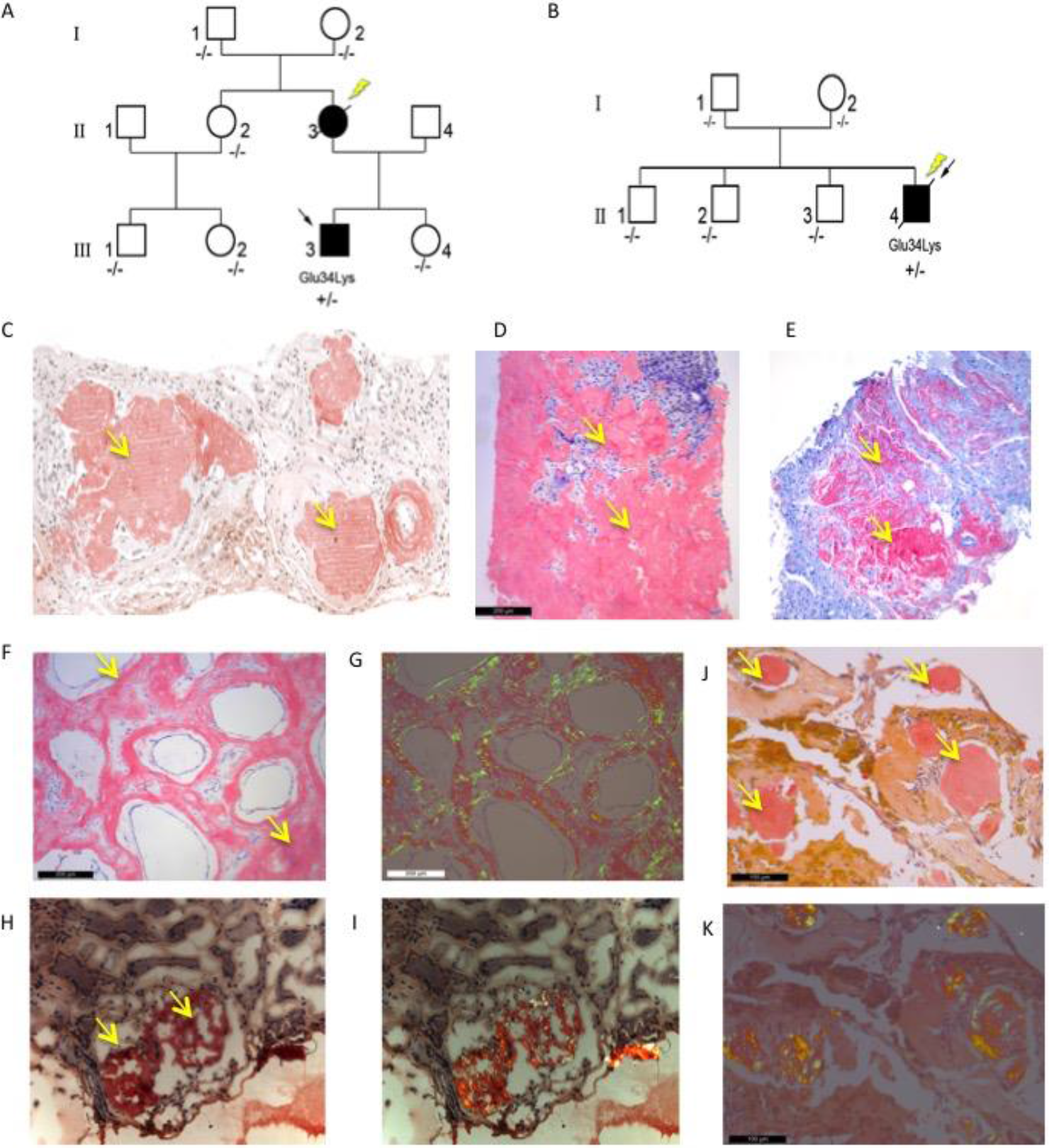
Anatomopathological features of AApoA-I associated with Glu34Lys variant. A. The pedigree of family A with the co-segregation of Glu34Lys. B. The pedigree of family B with the co-segregation of Glu34Lys. Black arrows indicate the probands. Glu34Lys/-designates the heterozygous mutation; −/− designates the absence of Glu34Lys on both alleles. Yellow arrows indicate the generation in which Glu34Lys has occurred *de novo*. C. Congo Red staining of kidney specimen derived from patient III.3 of family A revealing reddish-orange amyloid deposits (yellow arrows). D. Congo Red staining of liver derived from patient III.3 of family A revealing reddish-orange amyloid deposits (yellow arrows) E. Congo Red staining of cardiac amyloid deposits (yellow arrows) from patient II.2 of family A. F. Congo Red staining of amyloid deposits (yellow arrows) in testes from patient III.3 of family A. G. Congo Red staining of amyloid deposits in testes from patient III.3 of family A shows apple-green birefringence under polarized light. H. Congo Red staining of amyloid deposits (yellow arrows) in kidneys from patient II.4 of family B. I. Congo Red staining of amyloid deposits in kidneys from patient II.4 of family B shows apple-green birefringence under polarized light. J. Congo Red staining of amyloid deposits (yellow arrows) in retina from patient II.4 of family B. K. Congo Red staining of amyloid deposits in retina from patient II.4 of family B shows apple-green birefringence under polarized light.

The proband’s mother (II.3) had developed nephrotic syndrome and renal insufficiency at age 28, and kidney biopsy revealed amyloid in the glomerular compartment. Over the following four years, she developed histologically proven amyloidosis of the liver and colon, ocular manifestations, and left ventricular hypertrophy for which cardiac amyloidosis was histologically confirmed (Fig 1E). The medullogram and bone marrow biopsy were negative for an underlying lymphoproliferative plasma cell disorder. However, because of increased serum levels of free immunoglobulin light chain and the absence of familial history of amyloidosis, proband’s mother (II.3) was diagnosed with light-chain amyloidosis (AL), which is a relatively common sporadic systemic amyloidosis. As a treatment for AL, autologous bone marrow stem cell transplantation was performed. Unfortunately, the patient died of infection brought on by this treatment.

Family B (Fig 1B): A 39 year-old man (proband II.4) presented with a 20-year history of hypertension, infertility with severely decreased testosterone levels, chronic diarrhea, and a persistent subnephrotic range of proteinuria (2.01 g / 24 h) associated with renal and liver dysfunction. Neurological examination was normal, but electromyography revealed mild axonal sensory motor polyneuropathy. Cardiac amyloidosis was suggested by echocardiogram and electrocardiogram, but heart biopsy was not performed. Free immunoglobulin light chains were within a normal range, and immunofixation of serum or urine showed no evidence of monoclonal gammopathy. Tests of liver function showed high serum alkaline phosphatase (854 UI/I), high γ-glutamyl transpeptidase (594 UI/L), and moderately increased aspartate aminotransferase (69 U/L) and alanine aminotransferase (63 U/L). Abdominal ultrasound revealed hepatomegaly and splenomegaly, and abdominal tomodensitometry showed calcified adrenal glands. Kidney biopsy revealed amyloid deposits in glomeruli, and liver biopsy showed large amyloid nodules localized at portal tracts (Fig 1H and I). Although all tests for an underlying inflammatory or infectious condition were negative, the diagnosis of amyloid A amyloidosis secondary to urogenital tuberculosis was suspected on the basis of widespread abdomino-pelvic calcifications. Despite antituberculosis treatment, the renal and liver functions gradually deteriorated, with proteinuria reaching 8 g / 24 h, and the development of bleeding esophageal varices and hepatic encephalopathy. Ultimately, this patient died from renal and liver failure at age 49. He also suffered from an unexplained bilateral loss of vision. A retinal specimen was obtained during surgery for massive serous retinal detachment; histological analysis revealed amyloid deposits with typical apple-green birefringence upon Congo Red staining (Fig 1J and K).

### Genetic and proteomic analyses identify Glu34Lys apoA-I in amyloid

In family A, sequence analysis of coding regions of *APOA1* gene from the proband (III.3) revealed a heterozygous single-base substitution in exon 3 (c.172G>A) resulting in Glu to Lys substitution at amino acid position 34 of the mature protein, Glu34Lys (Fig 2A). While the proband’s mother (II.3) was dead and could not be tested, DNA from seven clinically asymptomatic family members was genotyped. Glu34Lys was not detected in the proband’s sister (III.4), father (II.4), aunt (II.2), and first cousins (III.1 and III.2). Importantly, Glu34Lys was absent from the DNA of both proband’s maternal grandparents (I.1 and I.2), suggesting strongly that Glu34Lys emerged *de novo* in the proband’s mother (II.3) (Fig 1A).

**Figure 2.**
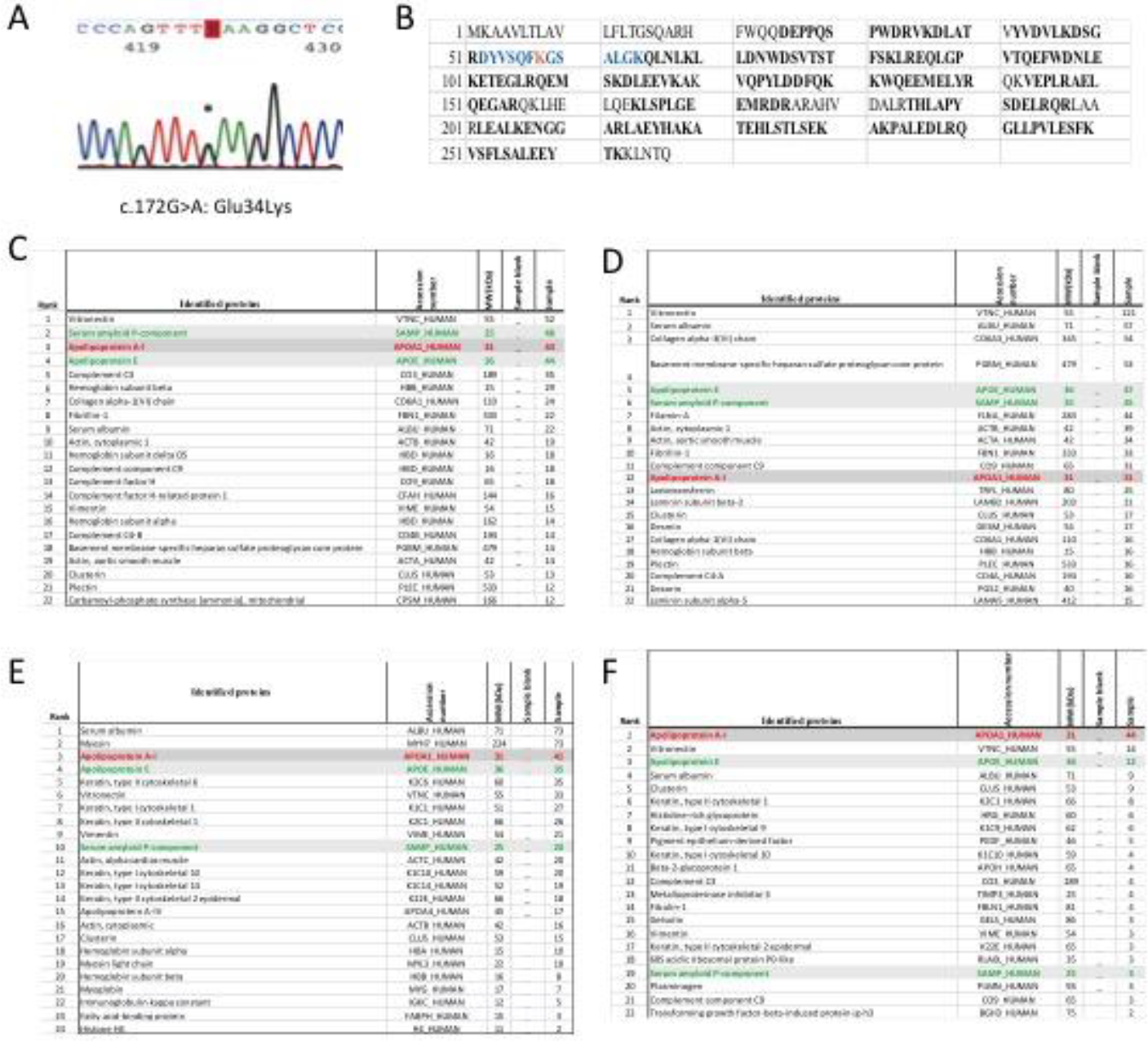
Genetic and mass spectrometric analyses of AApoAI Glu34Lys variant. A. Identification of the heterozygous c.172G>A mutation in exon 3 of *APOA1* by Sanger sequencing, resulting in Glu34Lys substitution. B. ApoA-I tryptic digestion followed by MS identified the mutation-containing peptide fragment (in blue) with lysine at position 58 of pre-apoA-I in red (nomenclature includes amino acids coding for the signal peptide, which corresponds to Lys34 of the mature protein). C. Liver amyloid proteome. D. Testis amyloid proteome. E. Retinal amyloid proteome. F. Heart amyloid proteome. All amyloid proteomes indicate the presence of apoA-I in fibrils, along with the “amyloid signature” proteins, apoE and serum amyloid P

In family B, Glu34Lys was also identified in proband II.4 in heterozygous state, but was absent from DNA samples of both his parents (I.1 and I.2). Consequently, Glu34Lys arose *de novo* in proband II.4 (Fig 1B).

To identify the fibrillar protein in amyloid, proteomic analysis of microdissected Congo-Red-positive deposits was performed using testicular and liver biopsies from proband III.3 of family A (Fig 2C and D), cardiac biopsy from proband II.3 of family A (Fig 2E), and the retinal specimen from proband II.4 of family B (Fig 2F). The results revealed apoA-I peptide fragments in amyloid deposits from all these organs, along with apoE and serum amyloid P, which are the “amyloid signature” proteins. Importantly, an apoA-I tryptic peptide containing Lys34 was detected in all four different tissues analyzed from the three Glu34Lys carriers, indicating amyloid deposition of Glu34Lys variant in these tissues (Fig 2B).

### Glu34Lys apoA-I is associated with a massive choroidal amyloid angiopathy

All three affected Glu34Lys carriers presented with a history of transient episodes of bilateral visual loss and metamorphopsia in their 30s. Ophthalmological investigations, which were available for proband III.3 of family A and proband II.4 from family B, showed similar clinical features. Fundus examination showed bilateral ovoid yellowish deposits at the midperipheral retina in the form of linear streaks and around the optic nerve head (Fig 3A). Retinal vessels were unremarkable, and no vitreous opacities were observed. Fundus autofluorescence revealed bilateral areas of autohypofluorescence in the peripapillary region, with a surrounding halo of hyperautofluorescence and with a hyperautofluorescent gravitational track from the optic disc to the inferior midperipheral retina (Fig 3B). Fluorescein angiography (FA) revealed bilateral staining around the optic discs, which was particularly prominent in the nasal peripapillary area. FA also showed a window effect of hyperfluorescent linear tracts of retinal pigment epithelium (RPE) atrophy, along with disseminated fluorescent spots, most probably at the level of RPE, predominating at the posterior pole (Fig 3C). Indocyanine Green angiography (ICGA) revealed hyperfluorescent linear radial streaks starting at the optic discs and following the course of large choroidal vessels at the posterior pole and at the midperipheral retina, as well as multifocal hyperfluorescent spots most probably located at the RPE level. Late-phase ICGA also revealed diffuse hypocyanescent dark lesions of variable size, some of which co-localized with yellowish deposits observed by funduscopy (Fig 3D and E). Therefore, the retinal phenotype observed in Glu34Lys carriers is characterized by a major choroidal angiopathy, without amyloid deposition in the vitreous, revealed at late-phase ICGA as hyperfluorescent streaks radiating from the optic discs and following the course of choroidal vessels with a diffuse “firework” pattern. To our knowledge, such a retinal phenotype associated with AApoAI has not been reported previously.

**Figure 3.**
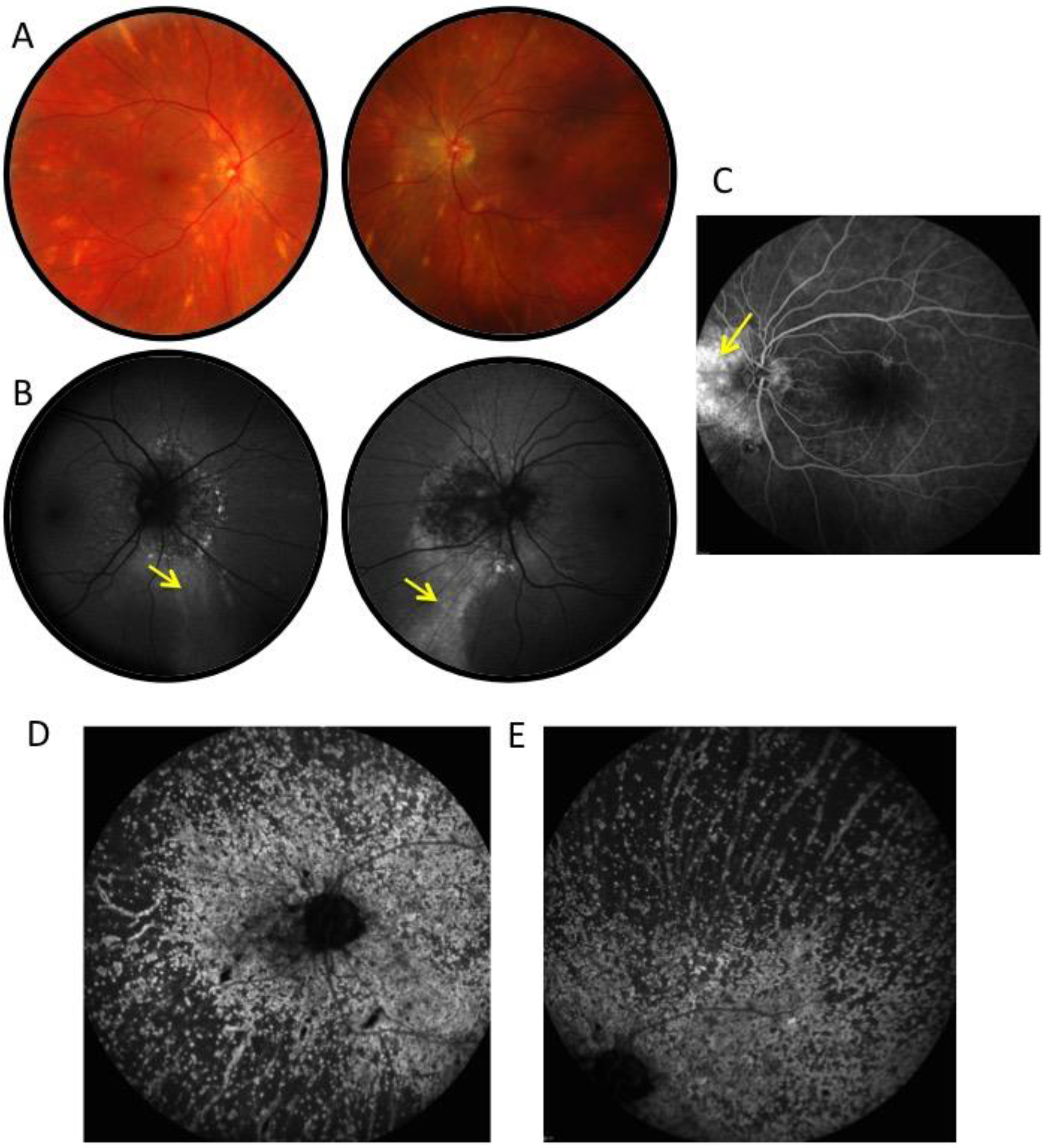
Ophthalmologic features of the choroidal/retinal AApoAI from proband III.3 of family A. A. Color fundus photograph of both eyes shows yellowish lesions around and irradiating from the optic disc. Some appear as black lesions (yellow arrows in B). B. FA shows the hyperautofluorescent gravitational track (yellow arrow). C. Autofluorescence shows intense staining of the nasal peripapillary area. D. ICGA shows diffuse hyperfluorescent spots predominantly around the optic nerve and linear hyperfluorescent lesions along the large choroidal vessels. E. ICGA shows diffuse hyperfluorescent spots and linear hyperfluorescent lesions along the large choroidal vessels at the posterior pole.

### Combined hepatorenal transplantation

After a multidisciplinary discussion that considered the extensive multisystem nature and the severity of the amyloid disease in patient II.4 from family B with fatal encephalopathy and blindness, a combined liver and kidney transplantation was proposed to proband III.3 from family A. The rationale for this dual transplantation was not only to offer a survival advantage by restoring normal liver and renal function, but also to slow the retinal damage, since the amyloid in choroidal vessels probably originated from hepatic production of variant apoA-I. At age 32, the patient underwent a liver and kidney transplantation from a cadaveric donor. No post-operative complications were observed, and there were no acute rejection episodes of either transplant. The patient reported a remarkable improvement of his general well-being and started exercising six months after the transplantation. Three years post-transplantation, both grafts were functioning very well, and recurring amyloid was not detected on renal biopsy. The patient did not develop any cardiac or neuropathic manifestations, and his visual symptoms have not progressed. Laboratory tests showed normal hepatic and renal functions. Hence, this dual transplantation was successful.

### Globular domains of variant apoA-I used as structural models

To elucidate the molecular basis for the highly amyloidogenic behaviour of Glu34Lys variant *in vivo*, we analysed its structure and stability *in vitro* and *in silico* using the lipid-free form, which is the protein precursor of amyloid (Das & Gursky, 2015). Unlike other AApoAI variants, Glu34Lys is the only known charge inversion variant and the only *APOA1* mutation located at the “top” of the four-helix bundle in lipid-free protein (Fig 4). This helical globular domain (residues 1-184) contains all known sites of AApoAI mutations, three out of four amyloidogenic segments, and the 9-11 kDa N-terminal fragments found in AApoAI deposits *in vivo* (Das *et al*, 2016). Therefore, the 2.2 Å resolution x-ray crystal structure of the globular domain of wild type (WT) apoA-I with the truncated C-terminal tail, Δ(185-234)WT (Fig 4) (Mei & Atkinson, 2011), provides an excellent starting model for understanding what makes apoA-I amyloidogenic (Gursky *et al*, 2012). This model was used in the current study.

**Figure 4.**
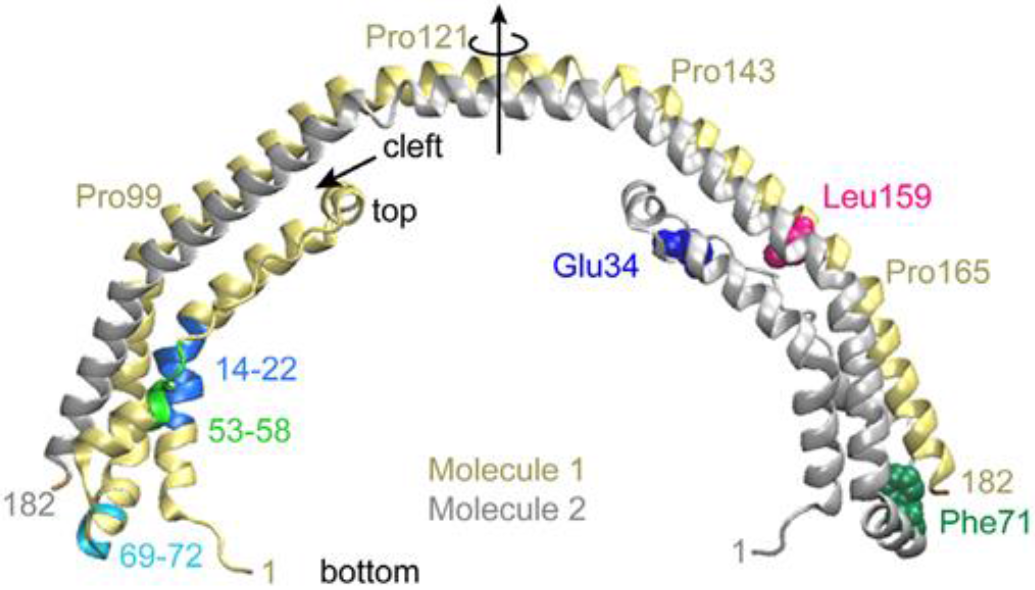
Atomic structure of the globular domain of lipid-free human apoA-I. The 2.2 Å resolution x-ray crystal structure of free e(184-243)WT (PDB ID: 3R2P) shows a crystallographic dimer comprised of two helix bundles. Dimer molecules 1 (yellow) and 2 (gray) are related by a two-fold symmetry axis (vertical arrow) that passes through the middle of the central linker (residues 121-142). Domain swapping around this flexible linker is thought to mediate monomer-to-dimer interconversion in apoA-I (Mei & Atkinson, 2011, Melchior *et al*, 2017). In the monomer, all segments in the four-helix bundle are from the same molecule. Top and bottom of the bundle are indicated. Short arrow points to the hydrophobic cleft between two pairs of helices, which is proposed to open upon lipid binding. Mutated side chains explored here, which are located in the hydrophobic core of the bundle, are shown in molecule 2 in a spherical representation: Glu34 (blue), Phe71 (green), Leu159 (pink). Amyloid hotspots are shown in molecule 1in residue segments 14-22 (blue), 53-58 (cyan) and 69-72 (light green) (Das *et al*, 2014, Das *et al*, 2016).

Here, we generated the recombinant globular domain of Glu34Lys apoA-I and explored its conformation, stability and dynamics. As controls, in addition to WT apoA-I, we chose two naturally occurring variants with very different biophysical properties and clinical presentations. Phe71Tyr is the most conservative AApoAI mutation that minimally perturbs the protein structure *in vitro* (Gursky *et al*, 2012; Das *et al*, 2016); *in vivo* it is associated with a very mild AApoAI restricted to amyloid in the palate without organ dysfunction. In contrast, Leu159Arg mutation, which greatly perturbs the protein structure *in vitro* (Das *et al*, 2016), is non-amyloidogenic *in vivo* but causes aberrant lipid metabolism and increased risk of cardiovascular disease (McManus *et al*, 2001; Sorci-Thomas *et al*, 2012).

Of all naturally occurring, full-length, recombinant apoA-I variants explored by our team (Das *et al*, 2014; Das *et al*, 2016), only Glu34Lys presented problems. This mutation led to a greatly decreased protein yield, and the protein precipitated after cleaving the tag, thereby precluding our attempts to obtain stable soluble full-length Glu34Lys apoA-I for biophysical studies. To bolster protein expression and solubility, the flexible hydrophobic C-terminal tail (residues 185-243) was truncated and the expression system was optimized to obtain ultrapure Δ(185-243) Glu34Lys as described in Methods. Previous biophysical studies of full-length WT, Phe71Tyr, and Leu159Arg variants (Das *et al*, 2016) provided useful controls for the current study of their globular domains.

Although the atomic structure of full-length free apoA-I is not available, strong evidence suggests that the C-terminal tail wraps around the 4-helix bundle and probably interacts weakly with the hydrophobic cleft between the two pairs of helices (Fig 4, short arrow) (Segrest *et al*, 2014; Mei *et al*, 2016). Since the secondary and tertiary structure of this 4-helix bundle in WT shows minimal changes upon the C-terminal truncation, Δ(185-243), the truncated WT provides a useful model for understanding the structure of full-length WT (Mei & Atkinson, 2011; Gursky, 2013; Melchior *et al*, 2017; Melchior *et al*, 2016). Here, we tested whether C-terminally truncated Phe71Tyr and Leu159Arg variants provided adequate structural models for their full-length counterparts, and determined how Glu34Lys, Phe71Tyr and Leu159Arg substitutions affect the structure and stability of the truncated protein.

### Glu34Lys destabilizes the globular domain and alters its aromatic residue packing

All proteins explored were pure and monodisperse in solution and formed dimers observed by SDS-PAGE and size-exclusion chromatography (Appendix Fig S1), similar to the crystallized construct (Mei & Atkinson, 2011). Far-UV CD spectra at 25 °C (Fig 5A) indicated an α-helical content ranging from 56±5% for l(185-243) Glu34Lys to 62±5% for Δ(185-243) WT, similar to 60±5% α-helix (or ~110 residues) reported previously for Δ(185-243) WT (Mei & Atkinson, 2011). Comparison with full-length WT, which showed 50±5% α-helix, including ~110 residues in the globular domain (Mei *et al*, 2016), suggested that this domain had very similar secondary structure in the full-length and C-terminally truncated WT. Like C-terminally truncated Glu34Lys protein, full-length Phe71Tyr and Leu159Arg proteins also showed a marginally significant helical loss upon single amino acid substitutions (Das *et al*, 2016). In summary, the helical content in the globular domain was perturbed slightly by the point substitutions in this domain, such as Glu34Lys, Phe71Tyr, and Leu159Arg, but did not significantly change upon C-terminal truncation.

Near-UV CD spectra were used to probe the aromatic residue packing in the globular domain. This domain contains all four tryptophans and five out of seven tyrosines in apoA-I, which dominate its near-UV CD spectrum. Importantly, Glu34Lys, Phe71Tyr and Leu159Arg variants showed distinctly different spectra indicating altered aromatic packing (Fig 5B). Further, globular domains and full-length forms of WT, Phe71Tyr, and Leu159Arg showed very similar near-UV CD spectra (Fig 5B compared with Fig 2C in (Das *et al*, 2016)). Moreover, near-UV CD of truncated and full-length recombinant WT resembled closely the spectrum of plasma lipid-free apoA-I (Mei & Atkinson, 2011; Das *et al*, 2016). These results show that Δ(185-243) truncation did not significantly change the aromatic side chain packing in the globular domain. However, this packing was altered by the point substitutions in this domain, such as Glu34Lys, Phe71Tyr or Leu159Arg.

Protein stability was probed by thermal denaturation wherein changes in the α-helical content were monitored by CD at 222 nm as described in the Supplementary Materials. All proteins showed a cooperative unfolding (Fig 5C) that was independent of the heating rate, indicating thermodynamic reversibility. The transition midpoint, measured with STD of 1.5 °C, was T_m_=7 °C in truncated WT and Phe71Tyr, significantly lower than that in full-length proteins (62 °C for WT and 60 °C for Phe71Tyr) (16). Truncated Glu34Lys showed T_m_=52 °C (Fig 5C). Truncated Leu159Arg showed the lowest T_m_=37 °C, which was 20 °C lower than that in the full-length Leu159Arg (Das *et al*, 2016). Truncated Leu159Arg also showed less cooperative unfolding evidenced by a decreased slope of the melting curve (Fig 5C, pink). These results are in excellent agreement with previous spectroscopic and hydrogen-deuterium exchange studies of full-length Leu159Arg, which suggested that Arg159 perturbs the core of the 4-helix bundle and splits it along the hydrophobic cleft, thus decreasing the protein’s stability and unfolding cooperativity and allowing the helix bundle to better sequester the C-terminal tail (Das *et al*, 2016). As a result, deletion of the C-terminal tail causes greater destabilization of the helix bundle in Leu159Arg than in Glu34Lys or other proteins.

The rank order of the protein stability emerging from the current study is WT≅Phe71Tyr>Glu34Lys>Leu159Arg (Fig 5C), in agreement with that observed in full-length proteins, WT≥Phe71Tyr>Leu159Arg (Das *et al*, 2016). This agreement suggests that, similar to their globular domains, full-length Glu34Lys is less stable than WT and Phe71Tyr, but more stable than Leu159Arg apoA-I. This result is consistent with the clinical finding that Glu34Lys mutation carriers have normal plasma levels of apoA-I and HDL, unlike Leu159Arg carriers whose plasma levels are reduced (McManus *et al*, 2001; Sorci-Thomas *et al*, 2012).

In summary, our biophysical studies revealed several new findings. Firstly, Glu34Lys charge inversion has minimal effect on the overall secondary structure in the globular domain of apoA-I (Fig 5A) but decreases its stability, as evidenced by a ~5 aC reduction in T_m_ (Fig 5C). Secondly, point substitutions in the globular domain, including Glu34Lys, Phe71Tyr and Leu159Arg, produce distinct near-UV CD spectra and hence, distinct aromatic side chain packing (Fig 5B and (Das *et al*, 2016)). Thirdly, despite its destabilizing effects, the C-terminal truncation has no significant effects on the secondary structure and aromatic packing in the globular domain of WT, Phe71Tyr, Leu159Arg (Fig 5A, B and C) and, by inference, Glu34Lys. These results validate the use of globular domains of these variant proteins as structural models for understanding their full-length counterparts. As such, they were employed in all molecular dynamics (MD) simulations, with initial coordinates drawn from the crystal structure of Δ(185-243)WT (Mei & Atkinson, 2011). CD data of these globular domains of WT, Glu34Lys, Phe71Tyr, and Leu159Arg (Fig 5) provided important experimental controls for these MD simulations.

### Mutations induce local unfolding and alter protein dynamics in MD simulations

To unravel early steps in the misfolding of apoA-I, we carried out MD simulations of the globular domains of WT, Glu34Lys, Phe71Tyr, and L159Arg proteins using the atomic structure of Δ(185-243)WT as a starting model. We simulated the monomeric apoA-I instead of the crystallographic dimer (Fig 4), as free apoA-I monomer is thought to form the precursor of amyloid; besides, the smaller monomer facilitated more extensive simulations. The monomer structure was obtained from the dimer via the domain swapping of one helical segment around the 2-fold axis (Fig 1 and 6B; see Supplementary Materials for details).

Firstly, the WT starting model was equilibrated at 300 K for 400 ns. Next, mutations were introduced, the structures were run at 300 K for 290 ns, and the last 250 ns were used for analysis. These room-temperature simulations were repeated thrice for each variant. The 4-helix bundle in all proteins remained stable (Appendix Fig S2). To diversify the conformational sampling, the dominant structure from the room-temperature simulations of each protein was heated to 500 K over 50 ps, followed by 20 ns of simulation at 500 K. The last 10 ns of these high-temperature simulations were used for analysis of helicity and correlation maps (see Supplementary Methods for details).

The 4-helix bundle in WT and Glu34Lys, Phe71Tyr and Leu159Arg proteins remained stable after high-temperature simulations but showed decreased secondary structure, mainly in residues 34-81 (Fig 6). Hence, the average α-helical content in the WT decreased from ~80% in the starting model to 60% in the final model. The latter agreed with the 61% α-helix content determined by CD spectroscopy in Δ(185-243) WT in solution (Fig 5) and with previous CD (Mei & Atkinson, 2011) and MD studies of this protein (Segrest *et al*, 2014; Melchior *et al*, 2016). Similarly, all variant proteins showed a decrease in their helical content by ~20% due to partial unfolding (Fig 6), in agreement with 56-62% α-helix observed in solution by far-UV CD (Fig 5).

**Figure 5.**
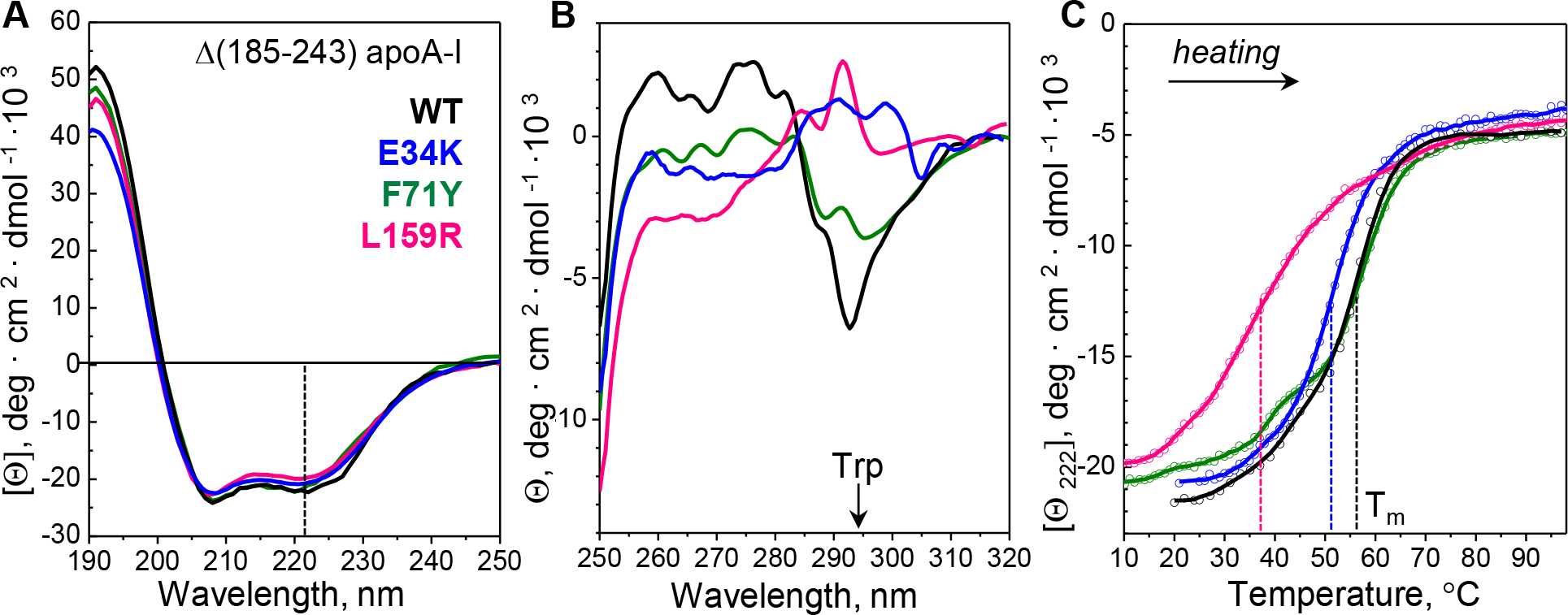
Conformation and stability of recombinant C-terminally truncated proteins. Globular domains (residues 1-184) of the WT (black), Glu34Lys (E34K, blue), Phe71Tyr (F71Y, green), and Leu159Arg (L159R, pink) were obtained and analyzed as described in Materials and Methods and Supplementary Materials. A. Far-UV CD spectra of the four proteins. Each spectrum represents 3-5 independent measurements. Helix content assessed from the CD signal at 222 nm (dashed line) ranged from 56±5% in Leu159Arg to 62±5% in WT. B. Near-UV CD spectra of the four proteins show large differences, particularly at wavelengths dominated by Trp (a peak centered at 295 nm). Each spectrum represents an average of three independent measurements with 5-point adjacent averaging. C. Melting data recorded by CD at 222 nm, Θ_222_(T), monitor α-helical unfolding during heating. Circles show raw data points. Dashed lines indicate melting temperatures, T_m_, corresponding to the first derivative maxima, dΘ_222_(T)/dT.

All proteins studied here showed full or partial unfolding in residues 34-81 (Fig 6) encompassing the second helical segment from the 4-helix bundle and adjacent hinge regions. Flexible secondary structure in this region is consistent with two large-scale motions, one around the “top hinge” near residue Gly39 and another around the “bottom hinge” containing Gly65-Pro66 and nearby groups; such hinge motions were proposed to mediate the helix-bundle opening during apoA-I binding to HDL (Mei & Atkinson, 2011; Gursky, 2013; Melchior *et al*, 2016). A kinked α-helix at the bottom of the bundle starting at Gln69 showed partial unfolding that was particularly pronounced in Phe71Tyr variant, probably due to the local perturbation around this mutation site (Fig 6).

**Figure 6.**
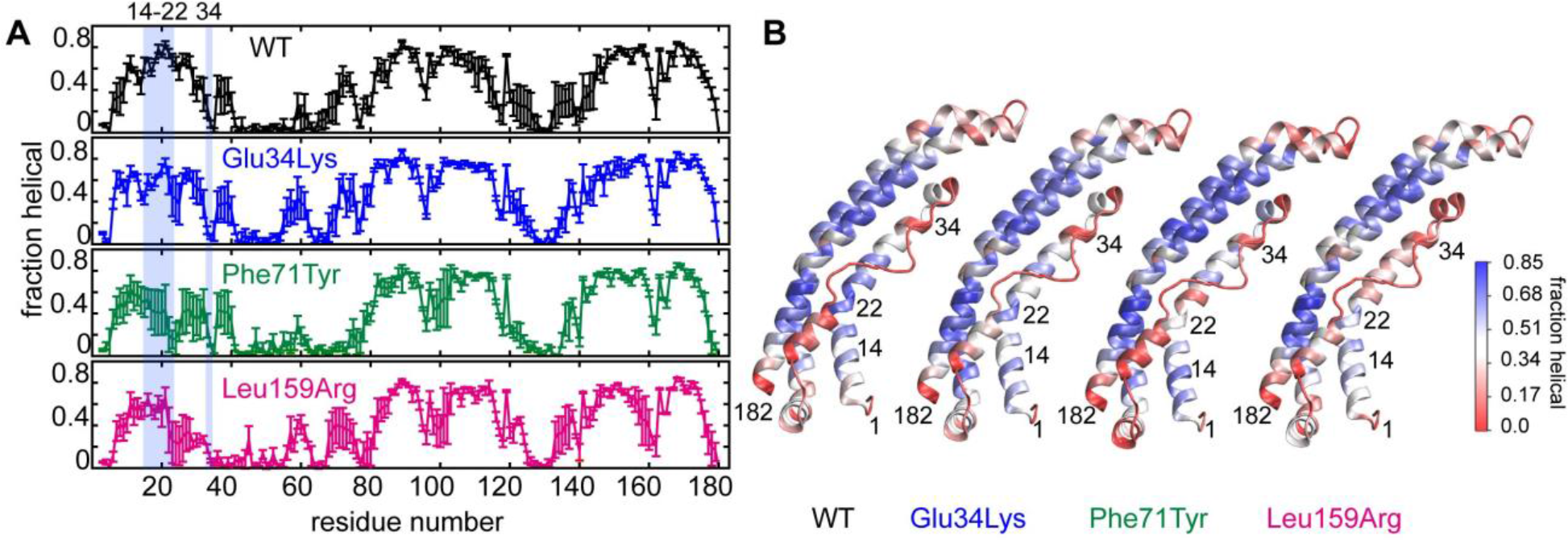
Variations in local helical conformation of WT, Glu34Lys, Phe71Tyr, and Leu159Arg proteins determined from high-temperature MD simulations. A. Helical fraction *versus* residue number in apoA-I variants. Standard errors of three replicates are shown by bars. B. The starting structure for each variant protein is colored based on the average helicity, from high to low helical fraction (blue to red), as indicated by the colored bar.

Despite these overall similarities, local mutation-specific differences were observed. This included the helical conformation at Tyr18 and downstream. Tyr18, which is located near a helical kink in the bundle, is in the middle of the major amyloid hotspot segment, Leu14-Leu22 (Fig 6B). Adhesive segments such as this are normally protected by their native packing from initiating protein misfolding. In WT, this segment is fully helical and thus protected, yet in all the three apoA-I variants this region shows an apparent helical loss (Fig 6A and B). We propose that such a mutation-induced loss of structural protection in this amyloid hotspot helps initiate apoA-I misfolding and aggregation. In fact, the structural protection observed in segment Leu14-Leu22 by hydrogen-deuterium exchange mass spectrometry (MS) declined in order WT>Phe71Tyr>Leu159Arg (Das *et al*, 2016). Hence, partial loss of helical structure in the major amyloid hotspot of Glu34Lys, Phe71Tyr and Leu159Arg, suggested by our MD simulations (Fig 6) and supported by experiment (*Das et al*, 2016), suggests a possible molecular basis for the amyloid formation by these proteins.

To determine how single amino acid substitutions influence concerted molecular motions in apoA-I, the pairwise correlation coefficients for C_α_ atoms were calculated for high-temperature structural ensembles. Fig 7A shows a correlation map for the WT; positively correlated regions move together (red), while negatively correlated regions move out of phase (blue). For each simulation, the average structure from three high-temperature replicates was calculated, and the absolute difference between Glu34Lys, Phe71Tyr and Leu159Arg variants and the WT was mapped to illustrate mutational effects on the molecular motion (Fig 7B, C and D). The difference map for Phe71Tyr showed few small areas where the variant protein deviated from the WT, while Glu34Lys and, especially, Leu159Arg showed much larger deviations (Fig 7B, C and D). Although the detailed interpretation of these maps was outside the scope of the current study, these results revealed that the effects of the point substitutions on the molecular motion were not localized to the mutation site but distributed across the protein, and progressively increased from Phe71Tyr to Glu34Lys to Leu159Arg (Fig 7B, C and D). A similar rank order was observed for the experimentally determined thermal stability, WT>Phe71Tyr>Glu34Lys>Leu159Arg (Fig 5C).

**Figure 7.**
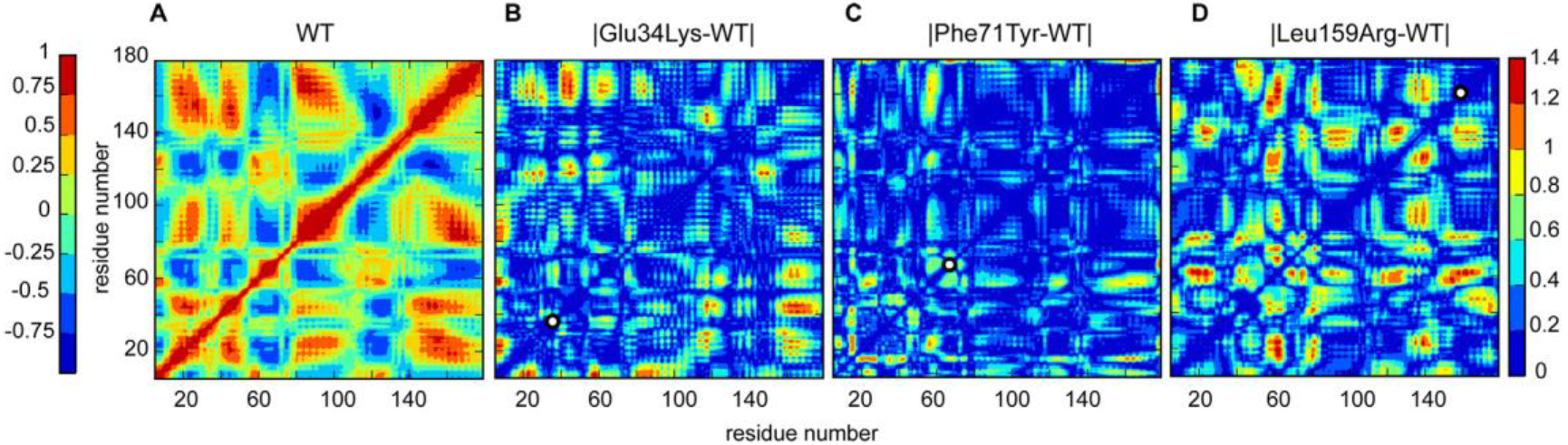
Correlation maps illustrate concerted molecular motions in WT, Glu34Lys, Phe71Tyr, and Leu159Arg proteins. The maps are calculated over three independent high-temperature MD simulations. A. Correlated molecular motions in WT depict protein groups that move either in concert (correlation coefficient 0 to +1, green to red) or out of phase (0 to −1, teal to blue), as indicated in the left color bar. B-D. Absolute difference between correlation maps of the three variants and WT apoA-I illustrates mutational effects on molecular motion. Groups whose relative motions remained invariant upon mutation are in blue and those whose correlated motions changed upon mutation are in warm colors (indicated in the right bar). Position of each apoA-I variant is marked with a white dot on the diagonal of each plot.

Together, our experimental and computational results consistently show that Glu34Lys charge inversion, which is much less conservative than Phe71Tyr substitution, is also more disruptive. Notably, the most disruptive mutation, Leu159Arg, is non-amyloidogenic *in vivo*. These results support the idea that structural destabilization alone is insufficient to make apoA-I amyloidogenic, and suggest that local protein conformation and dynamics in sensitive regions are important for amyloid formation (Das *et al*, 2016).

### AApoAI mutations induce a conformational shift in Trp72

Since Glu34Lys, Phe71Tyr and Leu159Arg variants have a distinctly different Trp side chain packing indicated by near-UV CD (Fig 5B), we analyzed MD trajectories to determine the conformational distribution of Trp in each protein. To depict local side chain motions, room-temperature trajectories were used. The dihedral angles χ_1_ and χ_2_ for each Trp were calculated at each step of the trajectory, and 2D histograms were obtained to compare the relative time spent in any given χ_1_, χ_2_ combination. ApoA-I contains four tryptophans (Trp8, Trp50, Trp72, and Trp108), one or more of which potentially contribute to the mutation-induced spectral changes in near-UV CD (Fig 5B). In the crystal structure, Trp8 and Trp72 are packed in the hydrophobic cluster at the “bottom” of the helix bundle, together with Phe71 and adjacent leucines; Trp50 forms a π-cation interaction with Lys23 in the middle of the bundle, and Trp108 together with Phe33 and Phe104 is packed in the “top” aromatic cluster (Mei & Atkinson, 2011).

MD simulations revealed significant mutational effects on the conformation of Trp72 (Fig 8). In WT, the vast majority of Trp72 rotamers clustered in a single region near χ_1_=295°, χ_2_=120°; this “closed” conformation represented the imidazole ring packed in the “bottom” hydrophobic cluster, similar to that seen in the crystal structure (Fig 8A). Although in mutant proteins this “closed” conformation remained predominant, its relative occupancy decreased and alternative conformations became populated (Fig 8B, C and D). In such “open” conformations, the imidazole ring of Trp72 pointed away from the hydrophobic cluster and into the solvent, thus allowing solvent entry in the bottom of the helix bundle. Such “open” conformations were highly populated in the two amyloidogenic mutants, Glu34Lys and Phe71Tyr, while the non-amyloidogenic Leu159Arg showed an intermediate population distribution between the amyloidogenic variants and the WT. Further, Trp8, which occupied a single conformation adjacent to Trp72 in the “bottom” aromatic cluster in WT, showed alternative conformations in Phe71Tyr and Leu159R but not in Glu34Lys. No alternative conformations were detected in Trp50 or Trp108 (Appendix Fig S3).

**Figure 8.**
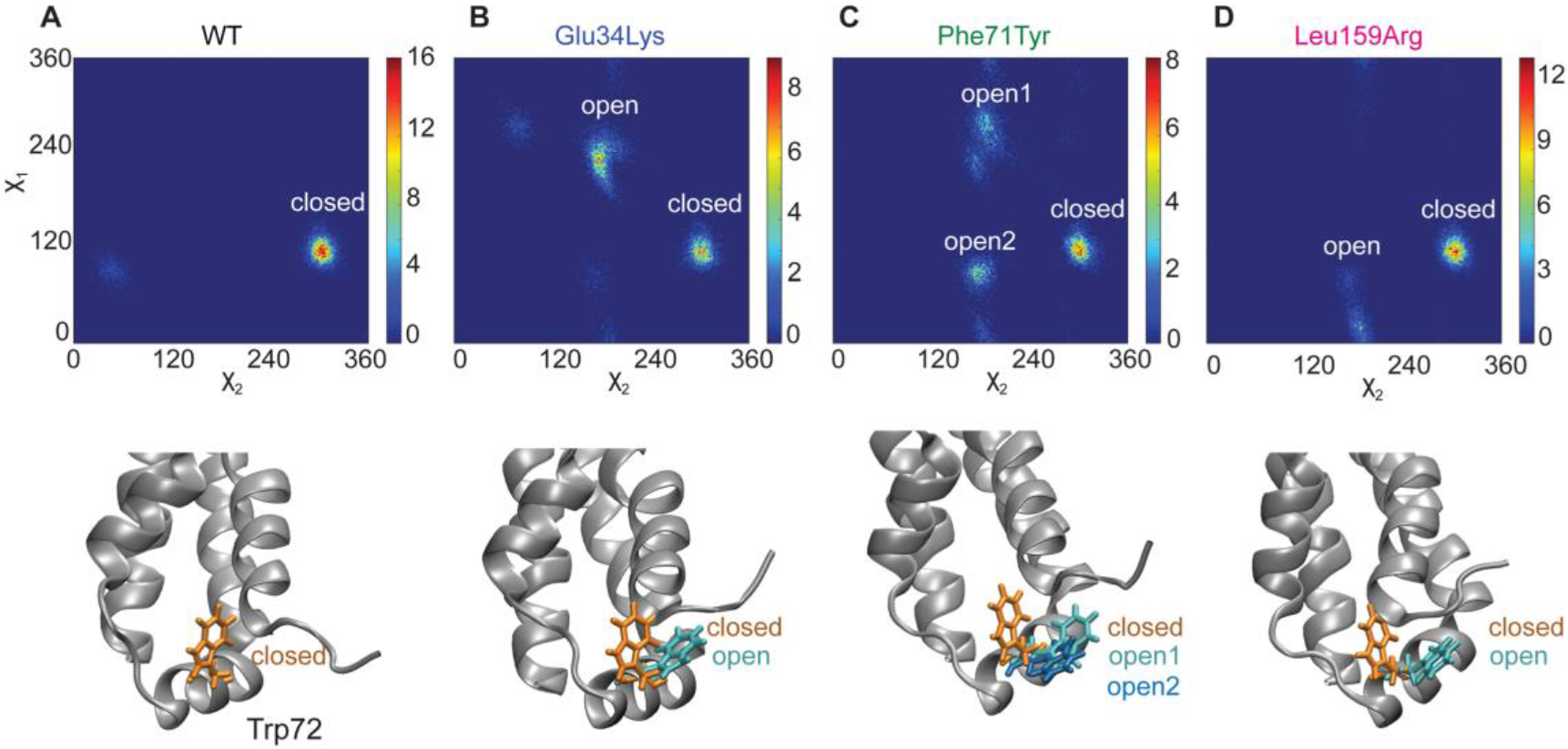
Conformational distribution of Trp72 side chain in WT and variant proteins. Probability distributions for χ_1,_ χ_2_ dihedral angles of Trp72 (top) and structural models depicting the corresponding conformations of Trp72 side chain (bottom) in proteins: A. WT, B. Glu34Lys, C. Phe71Tyr, D. Leu159Arg. One representative result of three independent room-temperature simulations is shown. The probability is proportional to the number of steps in the simulation trajectory. Least and most probable conformations are in blue and in warm colors, respectively, according to the color bars on the right. The color bars in different panels are different to illustrate full range of probabilities for different proteins. WT shows the sharpest peak corresponding to the “closed” Trp72 conformation. Glu34Lys, Phe71Tyr, and Leu159Arg proteins show additional peaks corresponding to “open” Trp72 conformations, which are depicted in bottom panels. Stick models show “closed” (orange) and “open” (cyan) orientations of Trp72 side chain.

In summary, MD simulations suggested that alternative conformations of Trp72 (Fig 8) and perhaps Trp8 (Appendix Fig S3) at the bottom of the 4-helix bundle were mainly responsible for the experimentally observed differences in the Trp packing among the variant proteins (Fig 5B). We propose that “open” conformations of Trp72, which were particularly highly populated in the two amyloidogenic variants, Glu34Lys and Phe71Tyr (Fig 8), could help initiate apoA-I misfolding by perturbing its structure at the bottom of the helix bundle. Such perturbations are expected to increase solvent exposure of the nearby Tyr18 and other residues from the major amyloid hotspot Leu14-Leu22, thus favoring protein misfolding and amyloid formation.

## Discussion

This study reports a new form of AApoAI that is caused by Glu34Lys variant and is associated with severely compromised kidney, liver, heart, testis, gastrointestinal tract, spleen and, unprecedentedly, the retinal/choroidal tissue. Our findings extend the phenotypic spectrum of this clinically heterogeneous disorder and highlight the complexity of its genotype-phenotype correlation, which is important for the timely diagnosis and treatment of this life-threatening disease.

Carriers of Glu34Lys described here developed uncommon ophthalmic manifestations, which correlated directly with biopsy-proven massive amyloid deposition and identification of variant apoA-I within amyloid fibrils from the retina. This complete anatomopathological concordance indicates that Glu34Lys variant directly participates in the retinal disease and is its causative agent. Therefore, our study provides the first detailed clinicopathological investigation of ophthalmic abnormalities in AApoAI and suggests that retinal amyloidosis should be considered in AApoAI cases with ophthalmic manifestations of undetermined origin. This idea is supported by the previous report of unexplained ophthalmic manifestations in one sporadic Glu34Lys AApoAI patient (Andeen *et al*, 2014) and in two additional AApoAI patients from another kindred carrying the c.280-288del in-frame mutation (Persey *et al*, 1998); these retinal abnormalities remained unexplored and their causal link with amyloidosis was not established. Taken together with the current study, these reports prompt us to propose to search for choroidal amyloid angiopathy in all suspected AApoAI patients to detect the onset of symptoms at the earliest possible stage, which is crucial for optimal therapeutic management and visual prognosis of the patients.

To-date, two sporadic cases of Glu34Lys AApoAI have been reported, both presenting with proteinuria and chronic kidney disease (Rowczenio *et al*, 2011; Andeen *et al*, 2014); one case also involved testicular and conjunctival amyloidosis (Andeen *et al*, 2014). A detailed anatomopathological study of a renal specimen has been performed for one Glu34Lys carrier, demonstrating unusual amyloid deposits in the glomeruli (Andeen *et al*, 2014). Clinical and pathological renal features of three additional Glu34Lys patients described in the current study definitively establish the causal role of Glu34Lys apoA-I in glomerular amyloid deposition. This new renal presentation contrasts with the typical tubulo-interstitial nephritis described previously for other apoA-I variants, which manifested as a slowly progressing, non-proteinuric renal failure with amyloid deposition restricted to the renal medulla (Gregorini *et al*, 2005; Gregorini *et al*, 2015). We conclude that AApoAI is characterized by distinct renal patterns of amyloid which depend on the nature of the apoA-I variant.

We also show that Glu34Lys variant causes early-onset male infertility due to massive amyloid deposition in testes. A similar clinical presentation was reported for many Leu75Pro carriers (Scalvini *et al*, 2007; Scalvini *et al*, 2008). Our study suggests that testicular involvement in AApoAI is more common than previously thought, leading to decreased male reproductive fitness. This strong negative selection factor is expected to reduce the number of apoA-I mutant alleles in the general population, unless the new mutational rate is high (Haase *et al*, 2012). Here we demonstrate neomutational events in the *APOAI* gene, thus providing the molecular explanation underlying apparently sporadic cases of AApoAI. This finding helps explain why AApoAI persists in the general population despite early-onset decline in male reproductive fitness (Haase *et al*, 2012). Moreover, sporadic clinical presentation suggests that the frequency of AApoAI is underestimated, which can result in misdiagnosis with the more common forms of acquired systemic amyloidosis caused by the deposition of Ig light chain or amyloid A. This misdiagnosis may lead to inappropriate treatments, including those described in the current study. Our findings will improve the timely diagnosis of AApoAI, which is very challenging yet is key to the patients’ prognosis and treatment.

In AApoAI caused by the Glu34Lys variant, the onset of first symptoms occurs as early as 15 years of age, with disease progression involving extensive amyloid deposition in patients in their 30s or 40s, which ultimately lead to one patient’s death at age 49. Currently, there is no targeted therapy for AApoAI. A favourable outcome of combined liver-kidney transplantation performed here for the first time in a 32-year-old man with Glu34Lys variant, which was beneficial as judged by the 3-year graft survival and visual prognosis, warrants the use of such a pre-emptive life-saving approach in cases of severe AApoAI.

In addition to expanding the clinical presentation spectrum and improving the diagnostics and treatment of AApoAI, the current study also advances our understanding of the molecular events involved in apoA-I misfolding and amyloid formation. Usually, amyloidogenic mutations in globular proteins are destabilizing, which is thought to augment protein misfolding. Our biophysical data clearly show that Glu34Lys follows this general trend. The rank order of stability for the globular domain of apoA-I established in the current study is WT≥Phe71Tyr>Glu34Lys>Leu159Arg. Notably, non-amyloidogenic Leu159Arg is the least stable variant explored so far. This paradoxical observation is consistent with our previous studies of full-length apoA-I, which provided a molecular explanation why Leu159Arg variant is rapidly cleared *in vivo* rather than accumulated in amyloid (Das *et al*, 2016). Together, our studies show that structural destabilization *per se*, which can shift the balance between protein accumulation and clearance, is insufficient for amyloid deposition *in vivo*. Consequently, factors other than the overall decrease in the protein structural stability must be involved in AApoAI.

The current study of Glu34Lys variant helps identify such factors. Firstly, low solubility of this full-length apoA-I variant as compared to those investigated in our previous work suggests that this charge inversion augments the protein aggregation. ApoA-I has a large fraction of charged residues that are non-randomly distributed in amphipathic α-helixes and apparently form extensive stabilizing salt bridge networks (Segrest *et al*, 2014). In the crystal structure of the lipid-free WT globular domain, Glu34 does not form salt bridges and is separated by ~8Å from the closest oppositely charged group, Lys45. Nevertheless, charge inversion in Glu34Lys, which is located in the hinge region at the top of the helix bundle, is expected to disrupt the overall electrostatics balance in this flexible region (Gursky *et al*, 2012). Charged residues are key to apoA-I hydration (Benjwal & Gursky, 2010) which determines protein solubility and interactions with other molecules. Hence, charge inversion such as Glu34Lys is expected to alter electrostatic interactions not only among apoA-I molecules, thus altering protein solubility, but also between apoA-I and other molecules. For example, negatively charged heparan sulphate proteoglycans are expected to interact more favorably with the more electropositive Glu34Lys variant, potentially leading to increased local concentration of the variant protein and augmenting its deposition in the extracellular matrix of tissues (Rosu *et al*, 2015).

Secondly, our results suggest that transient opening of Trp72 represents a previously unidentified early step in the misfolding pathway of amyloidogenic variants of apoA-I, such as Glu34Lys. Indeed, we demonstrated experimentally that all apoA-I variants explored in the current study have altered aromatic side chain packing in the globular domain. Consistent with this finding, MD simulations revealed that WT and Glu34Lys, Phe71Tyr, Leu159Arg apoA-I variants exist in a dynamic equilibrium between the “closed” and the “open” Trp72 conformations, and that the two AApoAI variants, Glu34Lys and Phe71Tyr, shift the equilibrium towards the “open” conformation. Such a shift is expected to facilitate water entry into the hydrophobic core of the helix bundle, thereby increasing solvent exposure of the major amyloid hotspot residues Leu14-Leu22 in apoA-I, which triggers protein misfolding. We posit that the molecular motions in apoA-I, such as the dynamic opening and closing of the Trp72 side chain, may extend to other apoA-I variants and influence their amyloidogenic behaviour *in vivo*.

Our findings exemplify how diverse amino acid substitutions at various protein sites alter protein conformation in sensitive regions, with potential implications for the protein function and disease. Indeed, amino acid substitutions in various apoA-I locations, including Glu34Lys (an “inside” mutation at the top of the helix bundle), Phe71Tyr (an “inside” mutation at its bottom), or Leu159Arg (an “outside” mutation near the middle), alter protein structure in similar regions, such as Trp72 at the bottom of the bundle. This phenomenon may extend to other apoA-I variants as well as to other globular proteins and their disease-causing variants (Lucato *et al*, 2017). Future studies will explore how the conformational equilibrium in these and other sensitive regions of apoA-I is influenced by factors such as the local pH, the cellular membranes and other tissue-specific factors relevant to the environment *in vivo*.

## Materials and Methods

This study was approved by the Ethics Committee of the Hospital of Rennes and Limoges and was performed according to the Declaration of Helsinki. Samples from the patients and their family members were obtained after their written informed consent. Molecular genetic and histological analyses of patient-derived samples and the MS analysis of amyloid deposits are described in the on-line Supplement. Recombinant proteins were cloned, expressed in E. coli and purified using His-tag affinity chromatography and FPLC; the protein purity and identity were confirmed by SDS PAGE and intact mass check using MS. Protein self-association was determined by FPLC, and protein secondary structure, stability, and aromatic side chain packing were characterized by far- and near-UV CD spectroscopy. MD simulations were performed in explicit solvent with 150 mM NaCl at 1 bar pressure, by using GROMACS 5.0 simulation package (Abraham *et al*, 2015) and CHARMM36 force field (Best *et al*, 2012). Methodological details are provided in the Supplement.

## Acknowledgments

We thank family members of the patients for their participation in this study. We thank Pr. Frank Bridoux (Service de Néphrologie, CHU de Poitiers, et Centre National de référence des amyloses AL et autres maladies de dépôts d’immunoglobulines monoclonales, 86021 Poitiers, France) and Brigitte Nedelec (INSERM, Institut Imagine, Université Paris Descartes, Sorbonne Paris Cité, France) for their assistance with anatomopathological analyses and figure preparation, respectively. We are indebted to Dr. Shobini Jayaraman (Boston University School of Medicine, USA) for invaluable help throughout this project, and to Christopher J. Wilson (Northeastern University, Boston, USA) for performing intact mass check of recombinant proteins. We thank Dr. Xiaohu Mei for help at early stages of protein expression.

## Author contributions

Study concept, design, and study supervision: SV, OG and JE-S; patient care coordination and clinical acquisition: JC-A, PR-R, TF, PR, AD, DS, AB; acquisition of data: MD, AP, AB, IM, JC-A, PR-R, TF, PR, AB, MC, FP, NR-L, JM-G, CB; analysis and interpretation of data: SV, OG, MD, AP and JE-S; drafting of the manuscript: SV, OG, JE-S and AP.

## The paper explained

### Problem

Amyloid diseases, or amyloidoses, affect millions of patients worldwide and represent a major public health challenge. In these diseases, a protein loses its native structure and deposits as insoluble cross-β-sheet rich fibrils in various organs, causing organ damage. Various monogenic forms of systemic amyloidoses caused by misfolding of different proteins often have overlapping clinical presentations, greatly complicating the diagnosis and treatment. One example is hereditary apolipoprotein A-I (apoA-I) amyloidosis (AApoAI), an incurable life-threatening disease resulting from heterozygous mutations in the *APOA1* gene. Improved knowledge of the clinical spectrum of AApoAI and the molecular triggers of protein misfolding is necessary to improve the diagnosis, optimize treatment, and develop much-needed rational therapeutic targets for this debilitating disorder.

### Results

We describe a hitherto unreported severe amyloidosis of the glomeruli, liver, heart, testes and gastrointestinal tract, along with unprecedented massive retinal amyloidosis discovered in two unrelated families. Genetic and proteomics analysis of various affected tissues showed that amyloid fibrils contained Glu34Lys apoA-I variant. Importantly, ophthalmic manifestations in these patients stemmed from fibril formation by this variant protein, revealing that the retina is a target tissue for AApoAI. Familial segregation of Glu34Lys indicated that these apparently sporadic cases stemmed from *de novo APOA1* gene mutation in both kindreds. WE report a successful combined kidney and renal transplantation performed as a life- and vision-saving measure for one young affected mutation carrier. Biophysical analysis of recombinant proteins showed that Gly34Lys substitution destabilized the globular domain of the lipid-free protein and altered its aromatic residue packing. Molecular dynamics simulation indicated local helical unfolding and revealed a previously unknown early step in the protein misfolding pathway. This step involved transient opening of the Trp72 side chain, which is expected to perturb native packing in amyloid hotspots, triggering protein misfolding.

### Impact

Our findings provide new mechanistic insights into AApoAI, improve the early diagnosis of this clinically heterogeneous genetic disease, and help patients’ management. The finding of a choroidal/retinal angiopathy in AApoAI not only offers new information about the biology of this disease in ocular tissues, but may also have profound therapeutic implications. In fact, this study prompts combined hepatorenal transplantation in the early stages of the disease to forestall severe retinal damage and vision loss. Finally, our results provide new insights into early steps in apoA-I misfolding, suggesting that a shift in the equilibrium between a protective closed conformation of Trp72 and an open conformation influences the amyloidogenic behaviour of the protein. Studies such as this may help guide future therapeutic targeting of AApoAI, which is currently unavailable.

